# Epigenetic Consequences of Hormonal Interactions between Opposite-sex Twin Fetuses

**DOI:** 10.1101/2020.06.09.141242

**Authors:** Siming Kong, Yong Peng, Wei Chen, Xinyi Ma, Yuan Wei, Yangyu Zhao, Rong Li, Liying Yan, Jie Qiao

**Affiliations:** Center for Reproductive Medicine, Department of Obstetrics and Gynecology, Peking University Third Hospital, Beijing 100191, China; National Clinical Research Center for Obstetrics and Gynecology, Beijing 100191, China; Key Laboratory of Assisted Reproduction (Peking University), Ministry of Education, Beijing 100191, China; Beijing Key Laboratory of Reproductive Endocrinology and Assisted Reproductive Technology, Beijing 100191, China; Beijing Advanced Innovation Center for Genomics, Beijing 100871, China; Peking-Tsinghua Center for Life Sciences, Peking University, Beijing 100871, China; Research Units of Comprehensive Diagnosis and Treatment of Oocyte Maturation Arrest; Academy for Advanced Interdisciplinary Studies, Peking University, Beijing 100871, China

## Abstract

In human opposite-sex twins, certain phenotypic traits of the female are affected negatively by testosterone transfer from the male, while the male may or may not be affected by the female *in utero*. However, no study was carried out to uncover the epigenetic basis of these effects. Here, we generated DNA methylation data from 54 newborn twins and histone modification data from 14 newborn twins, including female-female (FF), female-male (FM), and male-male (MM) newborn twins, to exclude the effects of postnatal environment and socialization, and investigated the epigenetic consequences of prenatal interactions between female and male gonadal hormones. We found that FM-Fs (female in FM twins) were distinguishable from FF twins by their DNA methylome, as were FM-Ms (male in FM twins) from MM twins. The correlation between genome-wide DNA methylation of females and males showed that FM-Fs, but not FFs, were closer to males from FM-Ms and MMs. Interestingly, the DNA methylomic differences between FM-Fs and FFs, but not those between FM-Ms and MMs, were linked to cognition and the nervous system. Meanwhile, FM-Ms and MMs, but not FM-Fs and FFs, displayed differential histone modification of H3K4me3, which were linked to immune responses. These findings provide epigenetic evidence for the twin testosterone transfer hypothesis and explain how prenatal hormone exposure is linked to reported and novel traits of childhood and adult through the epigenome in opposite-sex twins.

**Author Summary:** Prenatal exposure to testosterone may affect physiological, cognitive, and behavioral traits in females with male co-twins, while the males in opposite-sex twins present weak and inconsistent influences. In this study, we systematically investigated the hormonal interactions between opposite-sex twins in newborns from epigenetics including DNA methylation and histone modifications. We show that DNA methylome in FM-Fs (female in FM twins) was different from FF twins and their DNA methylomic differences were associated with cognition and the nervous system. We also suggest that FM-Ms (male in FM twins) were distinguishable from MM twins by their DNA methylome and FM-Ms *versus* MMs displayed differential histone modification of H3K4me3, which were linked to immune responses. Our study provides insight into the epigenetic explanation for hormonal influences between opposite-sex fetuses.

## Introduction

The incidence of spontaneous dizygotic twinning is between 1% and 4%[1], which is increasing due to advanced maternal ages and the application of assisted reproductive technologies[2, 3]. Approximately 40% of twins are opposite-sex (OS)[4]. Androgens, particularly testosterones, play important roles in early embryonic development[5]. Testosterone production in fetuses increases from 8 to 24 weeks of gestation and the maximal level occurs between 10 and 15 gestational weeks[6, 7]. It overlaps with the period of rapid brain development, as microglial cells colonize the cerebrum during 4 to 24 gestational weeks[8]. According to the twin testosterone transfer hypothesis, a female with a male twin is exposed to higher levels of prenatal testosterone[9]. Potential differences have been reported between OS and same-sex (SS) twins in physiological, cognitive, and behavioral traits[6]. The female OS twin may be more masculine, for example, leaning towards conservatism[10], breaking rules[11], having poorer performance in high school and college, being less likely to marry, having lower fertility, and earning less[12]. Female OS twins as children, but not as adults, also have larger total brain and cerebellum volumes[13]. Male OS twins, however, show weak and inconsistent evidence of behavioral and biological changes[12, 14, 15].

Although previous research has reported that a female exposed to prenatal testosterone from her brother exhibits specific behavioral and cognitive traits, the mechanism is still unclear, especially for epigenetic mechanisms. Animal studies have shown that testosterone triggers a DNA demethylation event in mouse embryonic neural stem cells and can also affect the global acetylation pattern of histone H3[16]. Cisternas *et al*[17] found that inhibiting DNA methylation in neonatal mice disrupted the testosterone-dependent masculinization of neurochemical phenotypes. Besides, human germline cells undergo global DNA demethylation from 7 to 19 weeks, coincident with testosterone production[18].

To study the underlying differences between OS twins, we generated epigenomic data from OS and SS twins, including DNA methylation and histone modifications, and explored potential influences of prenatal hormone exposure on them.

## Results

### Sex hormones have prenatal effects on the DNA methylome in opposite-sex twins

To systematically study the epigenetic differences between OS twins, we performed reduced-representation bisulfite sequencing on umbilical cord blood from 18 FF, 17 MM, and 19 FM twins. To determine whether FM-F and FM-M twins have unique DNA methylation patterns, we conducted hierarchical clustering on FM-F *versus* FF and FM-M *versus* MM twins according to their DNA methylation level at the top 3% CpG sites with the highest standard deviation. The results showed that FM-F and FF twins, as well as FM-M and MM twins, were clearly clustered into subgroups (Figs 1a-b), indicating that both FM-F and FM-M twins have features different from the corresponding SS twins.

**Fig 1.**
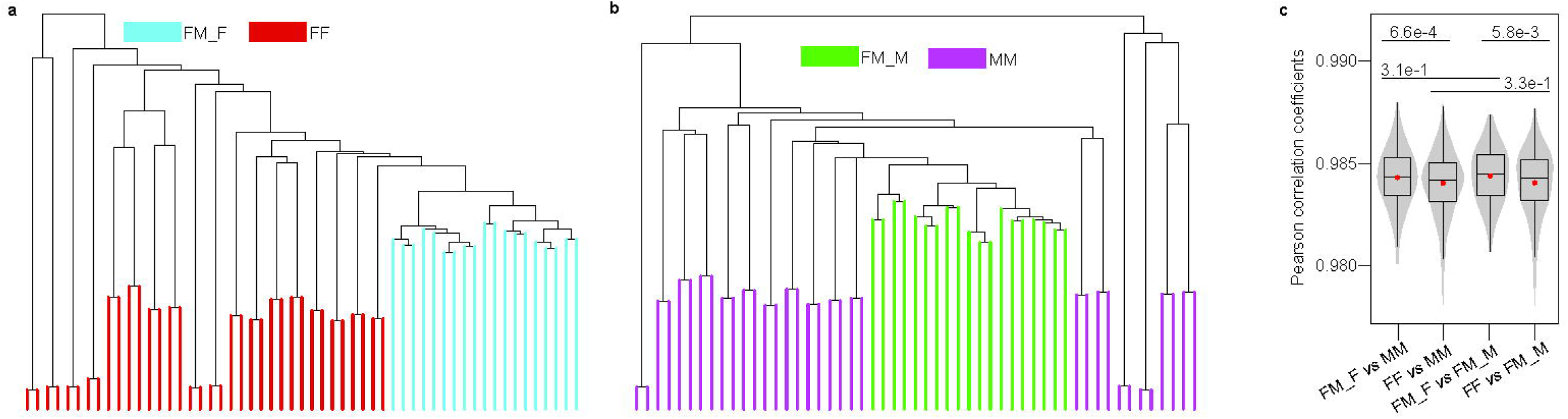
Characterization of genomewide DNA methylation in newborn twins. a-b. Hierarchical clustering for FM-Fs vs FFs (a), and FM-Ms vs MMs (b) based on DNA methylation level at the top 3% CpG sites with the highest standard deviation. c. Violin-box plots show the distribution of Pearson correlation coefficients between genomewide DNA methylations of any two samples from different two groups of twins. The red dots are the arithmetic means. The p-value between two groups was determined by Wilcoxon rank-sum test.

To further identify the distinction between FM-Fs and FM-Ms, we calculated the Pearson correlation coefficients between any two samples from different groups of the newborn twins. Interestingly, FM-Fs *versus* males (FM-Ms and MMs) had higher correlations than FFs *versus* males and FM-Fs were more strongly correlated with FM-Ms (Fig 1c). These results suggested that the DNA methylome of FM-Fs is not only more similar to their brothers, but also more similar to MMs comparing with FFs. Meanwhile, there were no significant differences between FM-Ms *versus* FM-Fs and MMs *versus* FM-Fs, as well as FM-Ms *versus* FFs and MMs *versus* FFs (Fig 1c). Furthermore, FM-Fs *versus* FM-Fs had the highest Pearson correlation coefficient among females in different types of twins while males in OS twins did not show this trend (S1 Fig). These results indicated that female OS twins are readily affected by their brothers and might express male characteristics, while male OS twins are also affected but have no female characteristics. Taken together, we can concluded that hormonal interactions between opposite-sex twin fetuses may induce different changes on the DNA methylome of FM-Fs and FM-Ms.

### Prenatal DNA methylomic changes are linked to the nervous system in females, but not males

To further study the potential biological functions of hormone-induced DNA methylomic changes, we identified differentially methylated cytosines (DMCs) with q-values <0.001 and methylation differences >20% in FM-Fs *versus* FFs, and FM-Ms *versus* MMs. The results showed that there were ∼1000 DMCs in different groups and only 10% (108) overlappsbetween hypo-DMCs of FM-Fs *versus* FFs and hypo-DMCs of FM-M *versus* MM, as well as 10% (108) overlapps between hyper-DMCs of FM-Fs *versus* FFs and hyper-DMCs of FM-M *versus* MM (S2a and S2b Figs). To further determine whether FM-Fs and FM-Ms induced the same changes, we performed cluster analysis based on their DNA methylation levels in all samples from the merged hyper-DMCs or hypo-DMCs. The results showed that the DNA methylation levels among female and male fetuses did not present consistent patterns in any cluster for both hyper-DMCs and hypo-DMCs (Fig 2a). Besides, the proportion of genes associated with overlapped DMCs was <20% (S2c Fig) and the genomic distances between DMCs in FM-Fs and FM-Ms were mostly >100 kb (S2d Fig). These results suggested that the influences of FM-Fs and FM-Ms might be different. To further explore the specific functions of FM-Fs and FM-Ms, we performed Gene Ontology analysis and found that the hyper-DMCs of FM-Fs *versus* FFs were mainly enriched in the processes of “nervous system” and “catabolic process” such as “neuropeptide catabolic process”, “hormone catabolic process”, and “collagen catabolic process”, and the hypo-DMCs of FM-Fs *versus* FFs were mainly enriched in “nervous system” such as “(negative) regulation of neuron projection regeneration and (negative regulation of axon regeneration)” and “integral component of postsynaptic membrane” (Figs 2b-c, S2e Fig). These results indicated that nervous system functions and catabolic processes in females in OS twins are mainly affected. However, the changes in DNA methylation in FM-Ms affected “RNA catabolic process” and “neuromuscular process controlling posture” (S2f and S2g Figs). Above all, these results indicated that the nervous system is probably affected in FM-Fs, but not FM-Ms, while both FM-Fs and FM-Ms may be vulnerable to catabolic influences.

**Fig 2.**
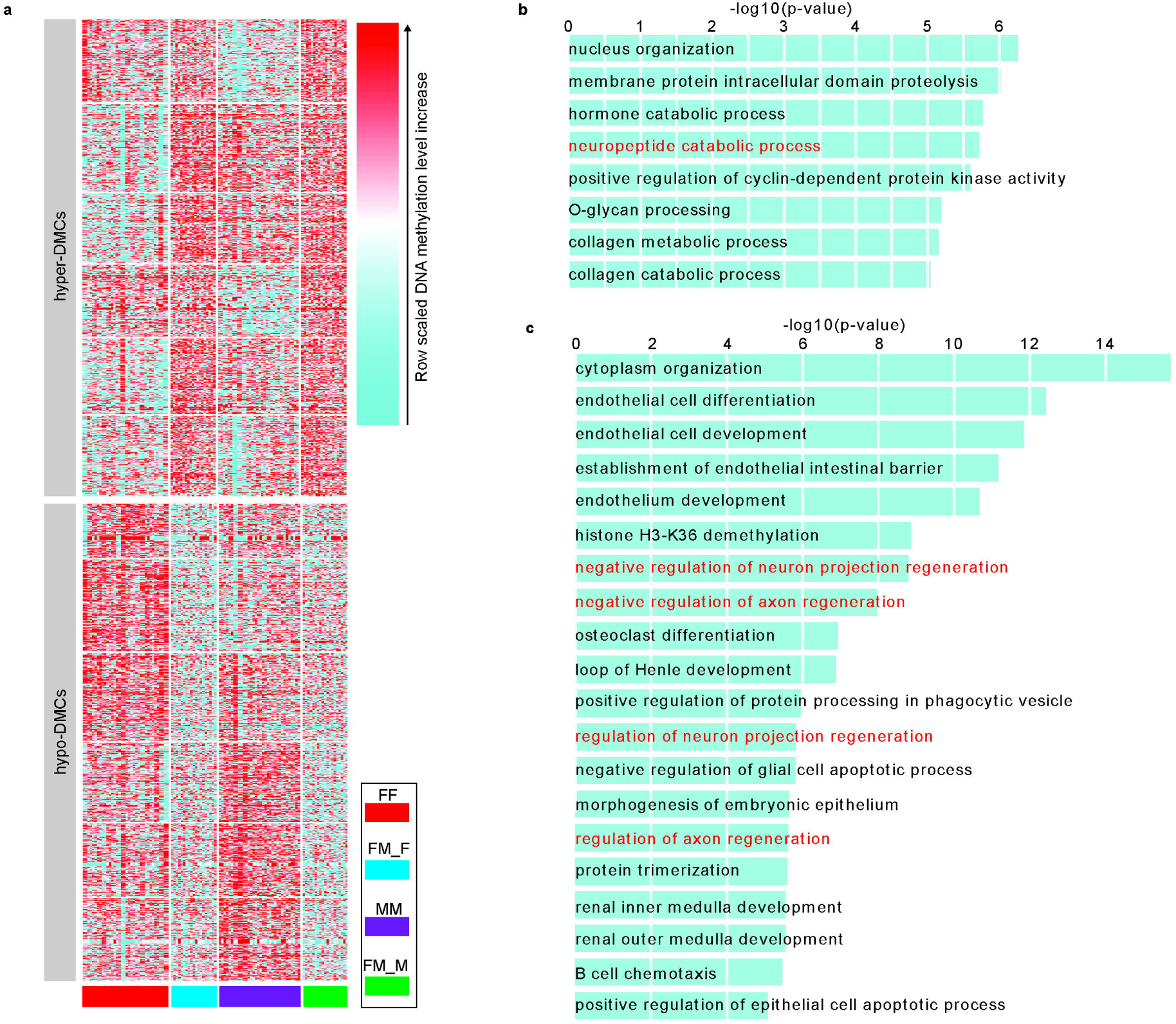
Differentially methylated cytosines (DMCs) in FM-Fs versus FFs, and FM-Ms versus MMs. a. K-means clustering for the all merged hyper- and hypo-DMCs. The hyper-DMCs or hypo-DMCs were merged if they have 1-bp common region at least, respectively. The hyper-DMRs (hypo-DMRs) were classified into six clusters by k-means clustering on row scaled DNA methylation level. b-c. Enrichment terms of gene ontology (biological process) with adjusted p-value < 10^−5^ were shown for the hyper-DMCs (b) and hypo-DMCs (c) of FM-Fs *vs* FFs. The GO terms associated with nervous system were highlighted by red.

### Prenatal histone modifications differ substantially between FM-Ms and MMs, but not FM-Fs and FFs

In addition to the DNA methylome, histone modifications can also be changed by the environment *in utero*[19, 20]. So, we also used chromatin immunoprecipitation sequencing (ChIP-seq) in core blood mononuclear cells from 7 FM, 4 FF, and 3 MM twins to examine changes in the histone modifications H3K4me1, H3K4me3, H3K27ac, and H3K27me3. Based on the genome-wide histone modifications, neither FM-Fs *versus* FFs nor the FM-Ms *versus* MMs could be clearly separated in hierarchical clustering analysis (S3a-S3d Figs). Differential peak analysis showed that >1000 GAIN H3K4me1 differential peaks (DPs) and GAIN H3K4me3 DPs were identified in FM-Ms *vs* MMs, while only a few GAIN/LOSS DPs were identified in FM-Fs *vs* FFs (S3e Fig). FM-Ms and MMs were clearly distinguished by GAIN DPs of H3K4me1 (Fig 3a) and functional enrichment analysis results showed that GAIN DPs of H3K4me1 between FM-M and MM fetuses were enriched in the processes of “gland morphogenesis”, “DNA replication”, and “transport”. In addition, FM-Ms and MMs were clearly separated by the GAIN DPs of H3K4me3 (Fig 3c), and their DPs were enriched in “immune response” (Fig 3d). Taken together, in contrast to DNA methylation, histone modifications may differ between FM-Ms and MMs, and are especially sensitive to the DPs of H3K4me1 and H3K4me3.

**Fig 3.**
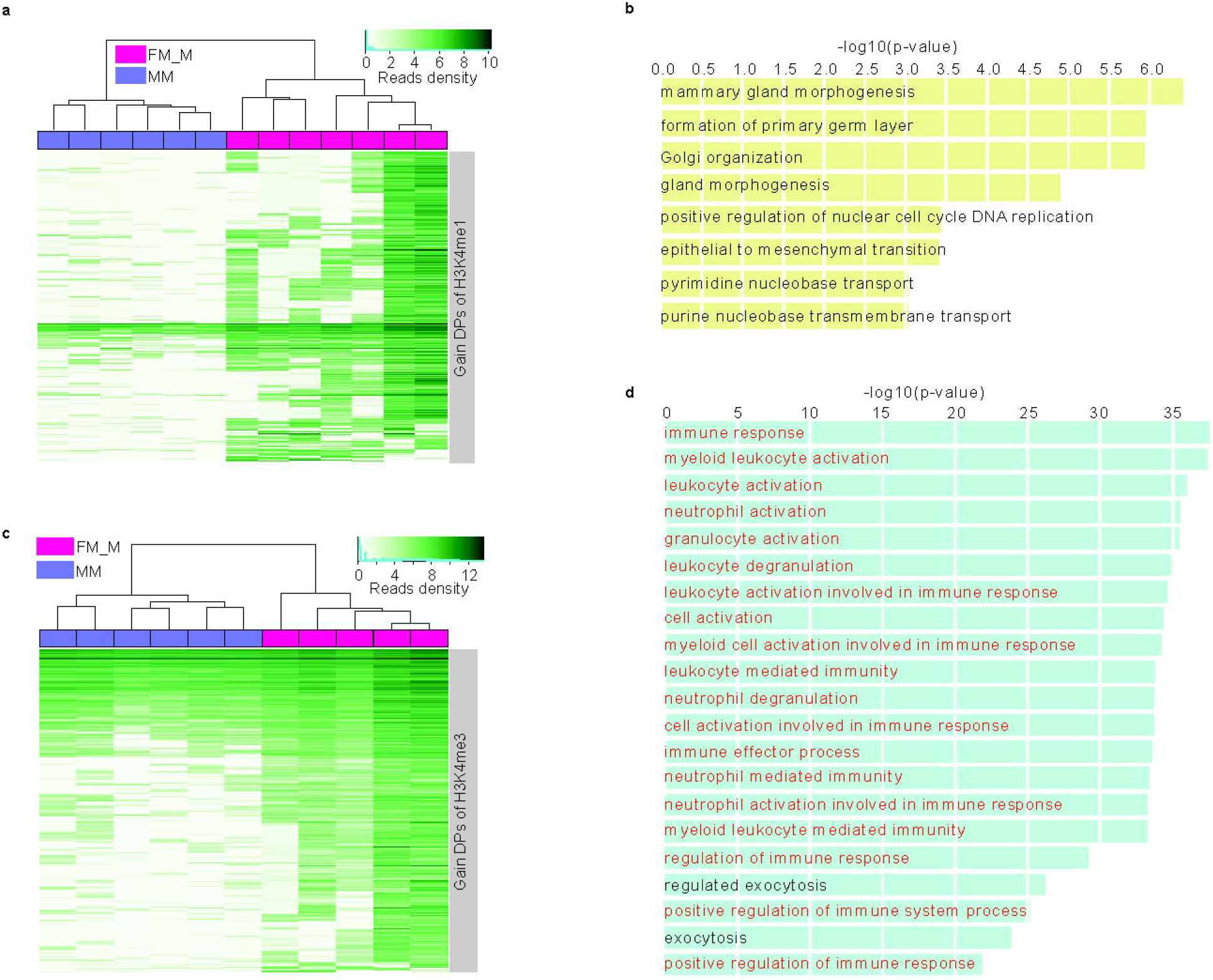
H3K4me1 and H3K4me3 differential peaks (DPs) in FM-Ms versus MMs. a. Hierarchical clustering for males in female-male (FM-M) and male-male (MM) twins based on H3K4me1 level at the DPs of H3K4me1. b. Enrichment terms of gene ontology (biological process) with adjusted p-value < 10-5 were shown for the 1034 GAIN DPs of H3K4me1. c. Hierarchical clustering for males in female-male (FM-M) and male-male (MM) twins based on H3K4me3 level at the DPs of H3K4me3. d. Enrichment terms of gene ontology (biological process) with adjusted p-value < 10-5 were shown for the 1643 GAIN DPs of H3K4me3. The GO terms associated with immune system were highlighted by red.

We further analyzed the relationship between DMCs and DPs, and the results showed almost no overlaps in the genomic regions while only a few DMC- and DP-associated genes (including GAIN H3K4me1 and GAIN H3K4me3) overlapped among all groups (S3f and S3g Figs). The genomic distances between DMCs and DPs (including GAIN H3K4me1 and GAIN H3K4me3) were mostly >100 kb (S3h Fig). The results implied that abnormal hormone effects have different influences on different levels of the epigenome of FM-Ms *in utero*.

## Discussion

Prenatal exposure to androgens may affect behavioral, perceptual, and cognitive traits in females with male co-twins; this supports the twin testosterone transfer hypothesis based on clinical phenomena[9]. However, the underlying mechanism is still unclear. In this study, we used >50 newborns to exclude postnatal environmental and social factors and focused on the epigenetic aspects of prenatal interactions. We systematically explored the differences in the genome-wide DNA methylation and histone modifications in OS and SS twins.

Our study first revealed that the DNA methylome in FM-Fs and FFs had different characteristics. FM-Fs *versus* FM-Ms showed stronger correlations than FFs *versus* FM-Ms, suggesting that FM-Fs are easily affected by their co-twin brothers. This is consistent with previous reports that a female who gestates with a male twin tends to be masculinized during development. An Australian cohort study involving 905 females with male co-twins and 3,964 female-female twin pairs reported that OS females have more socially-conservative views than SS females[10]. Studies of different age groups of OS females and SS females found that masculine characteristics occurred only in the group aged 17–28 years and the OS females scored higher on a rule-breaking subscale than SS females[11, 21]. Besides, OS females had a higher mean age at menarche than SS females, according to a study in Finland[22]. On the other hand, FM-Ms *versus* FM-Fs and MMs *versus* FM-Fs showed no significant differences, as did FM-Ms *versus* FFs and MMs *versus* FFs, FM-Ms are less influenced by their female co-twins. Previous cohort studies showed no significant differences between FM-Ms and MMs in cognitive traits such as sensation-seeking[23] and conservatism[10], but might have distinct features in autistic symptoms/hyperactivity disorder since FM-Ms had lower mean scores on the Social Responsiveness Scale than MMs[24].

DNA methylomic changes in FM-Fs, but not FM-Ms, were associated with the nervous system and this may play a crucial role in perceptual and cognitive activities. In a study of auditory system function, OS females had a lower frequency of spontaneous optoacoustic emissions than SS twins[25]. Females with male co-twins at 20 to 38 months of age tended to be somewhat weak in expressive vocabulary and scored lower on the MacArthur Communicative Developmental Inventory than SS females in two cohort studies [26, 27]. Females in OS twins also performed better than female-female twins in visuospatial cognition as assessed by the Mental Rotation Test[13, 28]. In an academic performance study in Denmark, FM-Fs obtained slightly lower ninth-grade test scores in mathematics than SS females, the opposite of that expected[29]. Males with female co-twins did not significantly differ in academic performance[29], auditory system function[25, 30], and visuospatial cognition[13]. Studies have also pointed out that female OS twins differ from SS females in neurobehavioral development in congenital adrenal hyperplasia (CAH), a genetic disorder caused by the overproduction of prenatal androgens. Such differences may involve activity interests, cognitive abilities, and personality[5, 31, 32].

Besides the nervous system, both FM-Fs and FM-Ms presented pronounced enrichment in catabolic terms. A Swedish cohort study of twins >60 years old showed that OS females had a moderately higher body mass index (BMI), body weight, and rate of dyslipidemia than female-female twins[33]. A comparative study in eight countries showed inconsistent BMI values in males with female co-twins[34]. Although no robust evidence has validated the influences on FM-Ms, they have numbers of DMCs similar to those of FM-Fs *versus* FFs and their potential risks need further attention.

Histone modification changes may regulate the chromatin status and gene expression[35]. Previous studies were mainly focused on the epigenetics of monozygotic twins[36]. Our study provides insights on histone modification differences in OS male twins. We found that the DPs of H3K4me1 and H3K4me3 may be a good marker to distinguish FM-Ms from MMs, but none of the four kinds of histone modification distinguished FM-Fs *versus* FFs. Interestingly, H3K4me3 in FM-Ms *versus* MMs might be related to immune responses. A Danish cohort study showed lower early-life mortality risks in OS boys than SS boys[29].

Our study focused on the epigenetic changes induced by sex interactions in OS newborns. Postnatal follow-up data are needed to further verify these changes. Furthermore, some traits may appear only after a long period of time, during which the corresponding phenotypes could be strongly impacted by environmental factors.

In conclusion, we provide an epigenetic basis to understanding the hormonal interactions in opposite-sex twins. Our results highlight that FM-Fs are affected by their male co-twins and the potential impact is mainly concentrated on the nervous system. Also, opposite-sex twins (both FM-Fs and FM-Ms) are prone to be affected by catabolic changes, and FM-Ms might be affected by immune responses. Long-term follow-up is necessary to provide more robust and extensive evidence.

## Methods

### Ethical approval

All blood samples were obtained after the participants gave written informed consent and were fully anonymized (each individual was given a code). The Reproductive Study Ethics Committee of Peking University Third Hospital approved the study (approved protocol no. 201752-044 in 2017/06/20). All relevant ethical regulations were followed.

### Study population and sample collection

OS and SS twins were born at the Third Hospital of Peking University; an accurate initial medical history confirmed that none suffered birth defects. A total of 56 families with twins were recruited and then divided into four main categories: FM-Fs, FM-Ms, FFs, and MMs. On the delivery day, ∼1 ml of umbilical cord blood was collected from each twin and processed immediately. Whole-blood aliquots of 300 µL were used for DNA extraction, and the remaining blood was stored at –80□ for backup. We also collected ∼20 mL blood and extracted cord blood mononuclear cells (CBMCs) as soon as possible.

### DNA extraction

The QIAmp DNA Blood Mini Kit (Qiagen, cat.: 51106) was used to extract genomic DNA (gDNA) from the stored whole blood samples.

### CBMC extraction and fixation

CBMCs were isolated by Ficoll density-gradient centrifugation (TBD, cat.: LDS1075).

Freshly-prepared 1% formaldehyde in 1× phosphate-buffered saline (PBS) was used to fix CMBCs for 10 min at room temperature. Glycine buffer (125 mmol/L) was added to quench the crosslinking reaction and the solution was incubated for another 5 min at room temperature. The fixed CBMC pellet was cryopreserved at –80□ after washing once with cold 1× PBS for subsequent ChIP-seq.

### Reduced Representation Bisulfite Sequencing (RRBS)

RRBS was performed following the published protocol with a few modifications^[37]^. In brief, ∼0.3% unmethylated lambda DNA (Thermo Scientific, cat.: SD0021) was spiked into 500 ng of high-quality gDNA, and the mixture was digested in MspI (Thermo Scientific, cat.: ER0541). The digested DNA was end-repaired and tailed with deoxyadenosine using Klenow Fragment exo- (Thermo Scientific, cat.: EP0422) after purification with Agencourt AMPure XP beads (Agencourt, cat.: A63881). Then NEBNext adapters (NEB, cat.: E7335) were used to ligate overnight with T4 DNA ligase at a high concentration (NEB, cat.: M0202M), and the Uracil-Specific Excision Reagent enzyme (NEB, cat.: M5505L) was used to digest the products.

Bisulfite conversion was conducted with a MethylCode Bisulfite Conversion Kit (Thermo Scientific, cat.: MECOV-50). The converted samples were size-selected by excising gel slices containing 160–700 bp DNA (2% TAE gel). A gel DNA recovery kit (Vistech, cat.: PC0313) was used to recover and then amplify the converted DNA fragments through PCR with a maximum of 12 cycles with Kapa HiFi U+ Master Mix (Kapa Biosystems, cat.: KK2801). Barcodes were introduced during this process.

The purified PCR-amplified products were quantified using Qubit dsDNA high-sensitivity dye in a Qubit 3.0 Fluorometer (Thermo Scientific, cat.: Q33216/Q32854).

A Fragment Analyzer™ Automated CE System (Analysis Kit: cat.: DNF-474-0500) was used to assess the fragment distributions and a Library Quant Kit for Illumina (NEB, cat.: E7630L) was used to measure molar concentrations. The final qualified libraries were sequenced using the PE150 strategy on an Illumina X Ten sequencer.

### ChIP-seq

ChIP-seq was performed as previously described^[38]^. Briefly, isolated nuclei were sonicated into fragments of 200–500 bp using an M220 Focused-ultrasonicator (Covaris) with the following parameters: burst, 200; cycle, 20%; and intensity, 8. Sheared chromatin in the supernatant was precleared by co-incubation with protein A Dynabeads (Invitrogen, cat.: 10002D) at 4°C for 1 h after centrifugation (13,000 rpm for 10 min at 4°C). The supernatant was diluted into 250 µL and transferred to low-binding Eppendorf (EP) tubes. The following antibodies were used for overnight incubation at 4□ with rotation: H3K4me3 (Millipore, cat.: 07-473), H3K27me3 (Abcam, cat.: ab6002), H3K9me3 (Abcam, cat.: ab8898), and H3K27ac (Abcam, cat.: ab4729). After washing with cold RIPA (10 mM, pH 7.6, HEPES, 1 mM EDTA, 4 mM LiCl, 1% NP-40, 0.1% N-lauryl sarcosine) and TEN (1 mM EDTA, 10 mmol/L, pH 8 Tris, 50 mM NaCl), 20 µg of Proteinase K (Qiagen, cat: 19131) were used to digest the immunoprecipitated pellets at 65°C for >6 h. DNA was purified using a MinElute PCR Purification Kit (Qiagen, cat.: 28006) and libraries were constructed using a NEXTflex™ ChIP-Seq Kit (Bioo Scientific, cat.: 5143-02). Fragments of 180–280 bp were size-selected using Agencourt AMPure XP beads (Beckman, cat.: A63881). DNA fragments were amplified by PCR with 12 cycles after tailing and ligating ChIP-seq adapters (Bioo Scientific, cat.: 514124). The final library was examined and sequenced as described for RRBS.

### RRBS data analysis

Before read mapping, adapters and low-quality bases in the reads were removed with Trim Galore to optimize the paired-end alignments. The kept reads were aligned to the *Homo sapiens* reference genome (human GRCh38/hg38) using Bismark (version 0.18.1) with the default parameters. The uniquely mapped reads with <2% mismatch were retained for further analyses. DMCs between any two groups were identified using methylKit with a q-value threshold of 0.001 and a mean methylation difference of 20%.

### ChIP-seq data analysis

To optimize the paired-end alignment, Trimmomatic was used for trimming before read mapping. The kept reads were aligned to the *Homo sapiens* reference genome (human GRCh38/hg38) using the Burrows-Wheeler Aligner with the default parameters in paired-end mode [39]. The non-redundant reads with ≤2% mismatches were retained for further analyses.

The MACS2 peak caller (version 2.1.0) with the parameter setting “--keep-dup=1, --broad” was used to identify peaks of histone modification. The R/Bioconductor package DiffBind was used [40], which uses DESeq2 or edgeR to identify DPs with q-values <0.05 and fold changes >2 for H3K4me1, H3K4me3, H3K27ac, and H3K27me3. The shared outputs of the DESeq2 and edgeR methods were taken as the final DPs.

### Clustering, Functional enrichment analysis, and Statistics

Hierarchical clustering was used for unsupervised clustering of the epigenomes of OS and SS samples with the R function ‘*hclust*’, in the R base package *stats*. K-means clustering was used for unsupervised clustering of DMCs using the R function *kcca* with the parameter “family=kccaFamily (which=NULL, dist=distCor)” in the flexclust package. To calculate distance, we used the *dist* function, and for similarity matrices we used the *cor* function of the *stats* package in R with the parameter “method=euclidean”. Potential biological functions of genomic regions, such as DMRs and DPs, were determined by GREAT with a p-value threshold of 10^−5^. For violin box-plots, the center dot indicates the average; the significance of differences between two groups was determined by the Wilcoxon rank sum test.

### Code availability

The bioinformatics pipelines and scripts used for our analysis are available at https://github.com/CTLife/Opposite-Sex-Twins_Epigenome

### Data availability

All the raw Data included in this study have been uploaded to the Sequence Read Archive database (https://trace.ncbi.nlm.nih.gov/Traces/home/) and are available for download via accession number GSE136849.

## Acknowledgments

This project was funded by the National Natural Science Foundation of China (81730038 and 81521002), the National Key Research and Development Program (2018YFC1004000, 2017YFA0103801, and 2017YFA0105001) and the Strategic Priority Research Program of the Chinese Academy of Sciences (XDA16020703).

## Author contributions

S.K., Y.P., W.C and L.Y. wrote the manuscript. W.C. and X.Y.M. collected the study materials and samples and patient data, performed the experiments. Y.P. developed the analysis methods and performed bioinformatics analysis. J.Q., L.Y. and W.C. developed the experimental concept and design. All the authors read and approved the final manuscript.

## Supporting information

**S1 Fig. Violin-box plots show the distribution of Pearson correlation coefficients between genomewide DNA methylations of any two samples from different two groups of twins**. The red dots are the arithmetic means. The p-value between two groups was determined by Wilcoxon rank-sum test.

**S2 Fig. Basic information of DMCs and GO enrichment**. a. Number of hyer- and hypo-DMCs for FM-Fs *vs* FFs, and FM-Ms *vs* MMs. b. Venn diagrams displayed the overlap among the DMCs. c. Venn diagrams displayed the overlap among the DMC-associated genes. d. Distribution of distances between two sets of DMCs. e. Enrichment terms of gene ontology (cellular components) with adjusted p-value < 10^−5^ were shown for the hyper-DMCs of FM-Fs *vs* FFs. f-g. Enrichment terms of gene ontology (biological process) with adjusted p-value < 10^−5^ were shown for the hyper-DMCs (f) and hypo-DMCs (g) of FM-Ms *vs* MMs. The GO terms associated with nervous system were highlighted by red.

**S3 Fig. Basic information of histone modifications**. a-d. Hierarchical clustering for females (left) and males (right) based on their H3K4me1 (a), H3K4me3 (b), H3K27ac (c), and H3K27me3 (d) level. e. Number of differential peaks of histone modifications. f. Venn diagrams displayed the overlap among the DMCs. g. Venn diagrams displayed the overlap among the DMC-associated genes. h. Distribution of distances between the two sets, DMCs and DPs.

