## Supplementary figures and images for "Epigenetic Consequences of Hormonal Interactions between Opposite-sex Twin Fetuses"

### S1_File.pdf

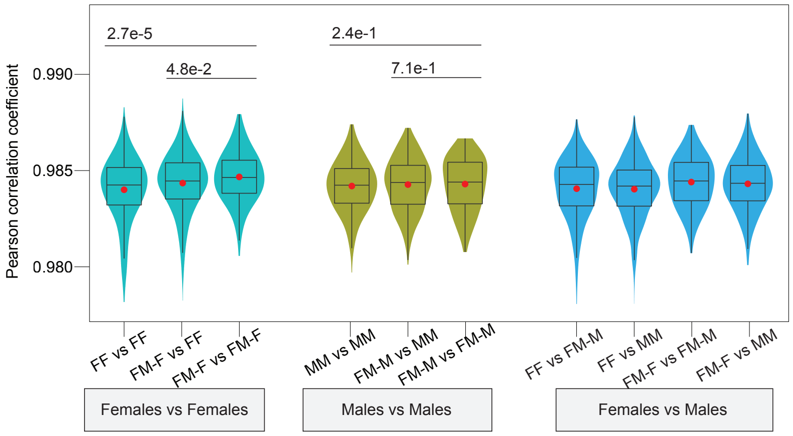

### S2_File.pdf

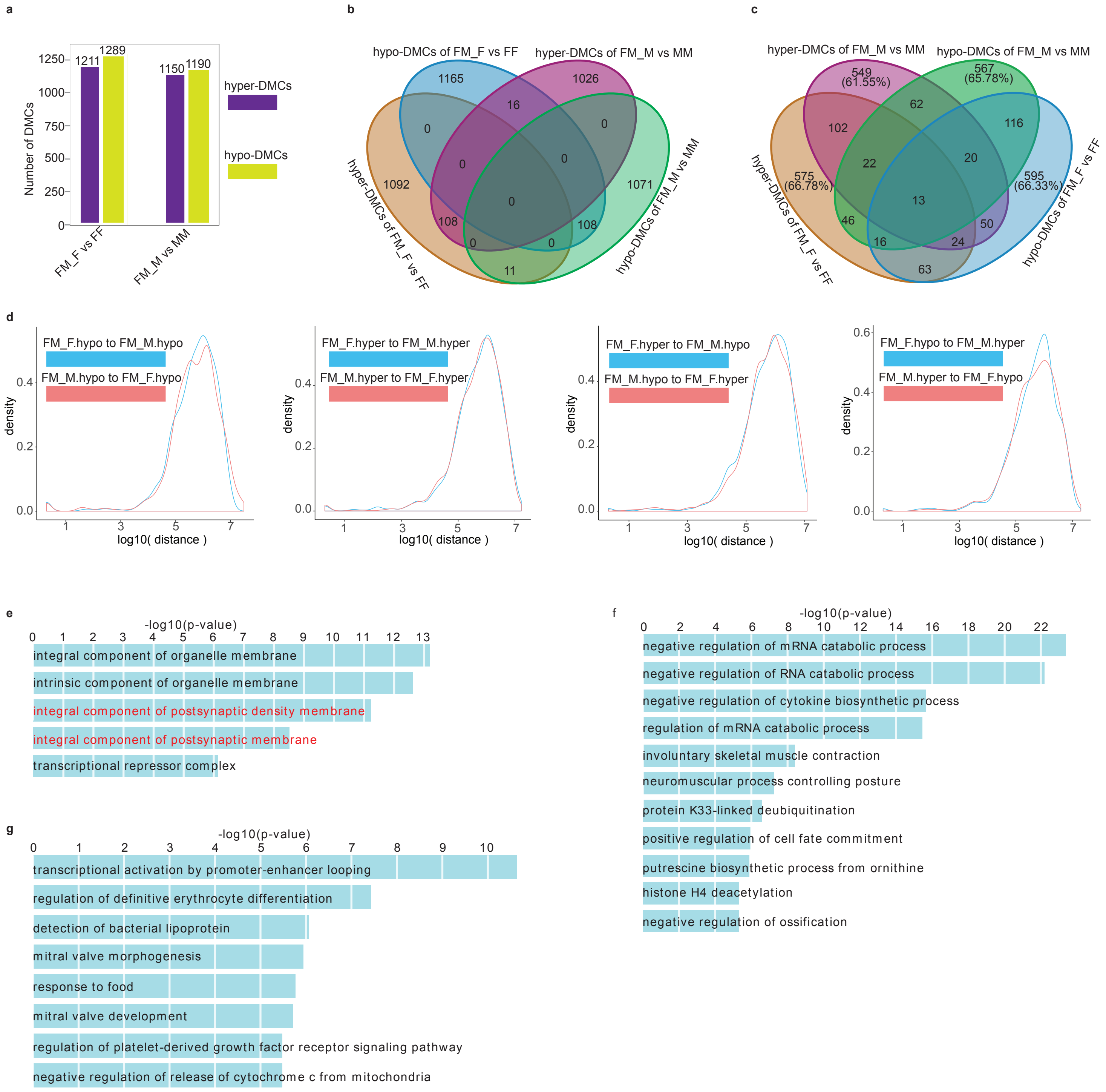

### S3_File.pdf

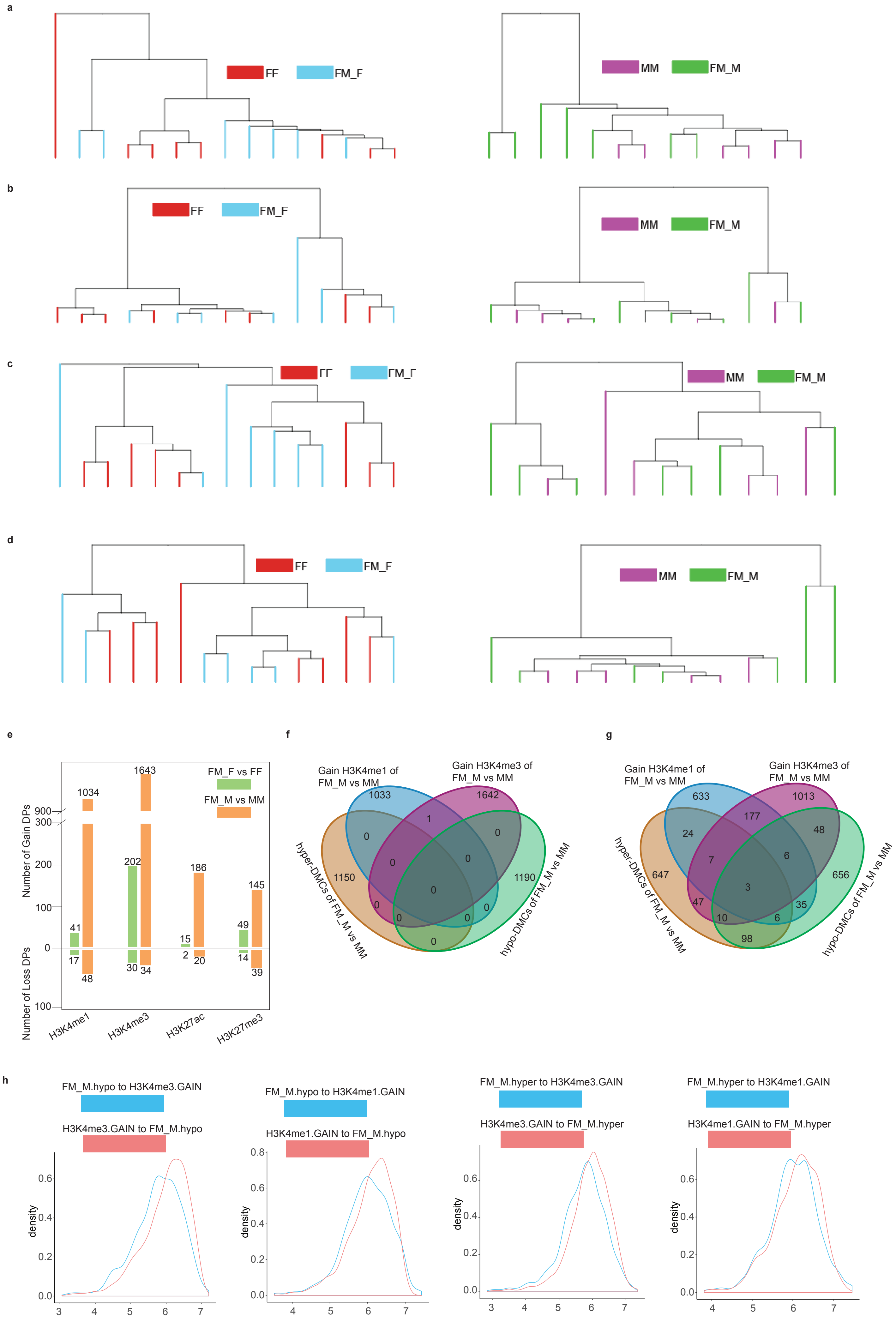
